# Optogenetic stimulation of nigral astrocytes is neuroprotective in a 6-OHDA model of neurodegeneration

**DOI:** 10.1101/2025.07.10.664169

**Authors:** Jessica L. McNeill, Ivan Trujillo-Pisanty, Stephanie Simard, Chase Groulx, Christopher A. Rudyk, Shawn Hayley, Maryam Faiz, Julianna Tomlinson, Michael Schlossmacher, Gianfilippo Coppola, Natalina Salmaso

## Abstract

**Highlights:**

- Optogenetic stimulation of nigral astrocytes attenuates motor deficits & Th+ cell loss in a 6-OHDA model of neurodegeneration
- Bulk RNA-seq analysis reveals optogenetic stimulation of nigral astrocytes induces early changes in microglia
- snRNA-seq shows 6-OHDA alone induces extensive gene expression changes across all cell populations within the SN
- Oligodendrocytes within the SNc express Th, which is upregulated with DA neuron loss

Parkinson’s disease is characterized by the loss of dopaminergic neurons in the substantia nigra. Glial-glial crosstalk is essential for maintaining the regional milieu, and appears to be particularly important in modulating neuroinflammation and many aspects of neurodegeneration. In particular, astrocytes are critical for maintaining dopamine neuronal integrity and survival, and astroglial dysfunction is prominent in Parkinson’s disease. As such, astrocytes represent a potentially critical therapeutic target in neurodegeneration. In this study, in vivo optogenetics were used to selectively stimulate astrocytes in the substantia nigra following a striatal 6-OHDA lesion. Remarkably, a single bout of optogenetic stimulation was sufficient to attenuate motor deficits and dopamine neuron loss induced by the neurotoxin. Furthermore, bulk RNA-seq and snRNA-seq analysis of the substantia nigra revealed extensive changes in both microglia and oligodendrocytes, suggesting that the neuroprotective effects of stimulating astrocytes may be mediated through alterations in glia-glia crosstalk. Altogether, this work demonstrates the importance of understanding glia-glia interactions in neurodegeneration.

## Introduction

Parkinson’s disease (PD) is a highly prevalent age-dependent neurodegenerative disease, characterized by the loss of dopamine (DA) neurons in the substantia nigra (SN) pars compacta and widespread Lewy pathology, largely comprised of misfolded forms of the α-synuclein protein.^1^ While the majority of studies have focused on nigral DA neurons to delineate the pathophysiological changes responsible for PD, it is increasingly evident that glial cells such as astrocytes, microglia and oligodendrocytes are critically involved in maintaining DA cell integrity.^2,3,4,5,6,7,8,9,10,11,12,13,14,15,16,17^

Astrocytes constitute an abundant and diverse group of macroglial cells, with extensive morphological, physiological and functional heterogeneity.^18,19,20,21,22,23^ They are critical components of the synapse: astrocytes release gliotransmitters, such as ATP, and modulate neuronal activity through the homeostatic regulation of ions and neurotransmitters (e.g. glutamate).^18^ Trophic support is also supplied by astrocytes through their release of FGF2 and GDNF growth factors that promote survival, development and plasticity of neighboring neurons. Astrocytes further promote neuronal survival through their antioxidant activity, as they are well known producers of glutathione and other antioxidant compounds.^18^ Importantly, astrocytes also play a critical role in the response to injury and disease through morphological, physiological and functional changes in a process referred to as reactive astrogliosis.^18^ Astrocytes can also act together with microglia to modulate neuroinflammatory cascades, being especially responsive to cytokines and participating in their release. For instance, the release of Il-1α, TNF-α and the immune system complement pathway protein, C1q, from activated microglia is sufficient to induce astrocyte reactivity and neurotoxicity in animal models.^24^ These pro-inflammatory factors are elevated in post-mortem brains of patients with PD and several parkinsonian models.^24,25,26,27,28,29,30,31,32,33,34,35,36,37,38,39,40,41^ Indeed, microglial-astroglial crosstalk seems critical to many aspects of neurodegeneration and recent studies have begun to also implicate the other primary glial type; namely, oligodendrocytes, in PD-like pathology. In fact, ferroptosis-induced DA neurodegeneration was reported to be mediated by oligodendroglial dysfunction, which, ultimately resulted in dysregulated astrocyte FGF signaling.^14,42^ Hence, it is reasonable to suggest that crosstalk between astrocytes, oligodendrocytes and microglia may be a crucial factor in the outcome following injury or disease.

Neurodegenerative disorders such as Parkinson’s disease are characterized by extensive changes in astrocyte functioning.^24,43,44,45,46,47,48,49,50,51,52,53,54^ For example, mutations in genes commonly associated with PD, such as LRRK2, DJ-1, PARK2 and GBA are known to alter lysosomal and mitochondrial function and impair glutamate uptake resulting in altered astrocyte inflammatory responses and reduce their antioxidant activity.^55^ The accumulation of α-synuclein is not only found in neurons, but also in in astrocytes in post-mortem tissue from patients with PD.^51,56,57,58,59^ Furthermore, genetic driven expression of the A53T mutant form of α-synuclein selectively in astrocytes was sufficient to cause an astrogliosis response that gave rise to motor deficits and midbrain dopaminergic cell death.^60^ This suggests that neuronal dysfunction and degeneration represents a failure of astrocytes to do their “job.” Consequently, taking the novel approach of targeting astrocytes could confer neuroprotective outcomes. Indeed, optogenetic stimulation of astrocytes *in vitro* has been shown to protect dopaminergic neurons from exposure to neurotoxins, and *in vivo* may promote regenerative effects of stem cells transplanted in the SN.^61^ In this study, we aimed to evaluate the potential therapeutic effects of *in vivo* optogenetic stimulation of astrocytes within the context of a neurotoxin 6-hydroxydopamine (6-OHDA) model of PD. We demonstrate that a single bout of nigral astrocyte optogenetic stimulation is sufficient to restore motor functioning and attenuate Th+ dopaminergic cell loss induced by 6-OHDA. Interestingly, bulk-RNA sequencing and single-nuclei RNA sequencing of the SN revealed extensive changes in microglia- and oligodendrocyte-related genes at 1- and 3-weeks post-lesion, respectively. Altogether, this work suggests that astrocytes are essential regulators of neurodegenerative processes and may be a critical target for therapeutic intervention in neurodegenerative disorders, like PD, in part, through changes in glia-glia interactions and crosstalk.

## Results

### Astroglial-specific Expression of AAV

In order to selectively stimulate nigral astrocytes *via* optogenetics, an AAV5-GFAP(0.7)-hChR2(C128S/D156A)-EYFP was employed. We first confirmed that AAV5-GFAP(0.7)-hChR2(C128S/D156A)-EYFP expression was restricted to astrocytes. Briefly, rats were unilaterally injected with 1.0 μl of the AAV into the SN (see *Methods*), and subsequently euthanized 5 weeks later. Immunohistochemical analysis of GFP, GFAP and S100β revealed that all GFP+ve cells overlapped with at least one astrocyte marker, and all cells showed astrocyte-like morphology, indicating specificity of viral expression to astrocytes (See Supplemental Figure 1).

### Nigral astrocyte stimulation reduces motor deficits

To determine the potential neuroprotective effects of optogenetic stimulation of nigral astrocytes, we employed a unilateral 6-OHDA model of neurodegeneration. This involved direct striatal injection to induce a relatively slow, retrograde degeneration of DA neurons that typically shows behavioral impairments by 3-weeks.^62^ To assess impairments, rats were tested on the accelerating rotarod, DigiGait and apomorphine-induced rotations test. Behavioral testing began 1-week or 3-weeks post-lesion (or saline control): the first day of testing consisted of the accelerated rotarod followed by gait analysis on the DigiGait system and two days later, rats were tested on the apomorphine-induced rotations test (refer to Figure 1A, B for the experimental overview and timeline).

**Figure 1.**
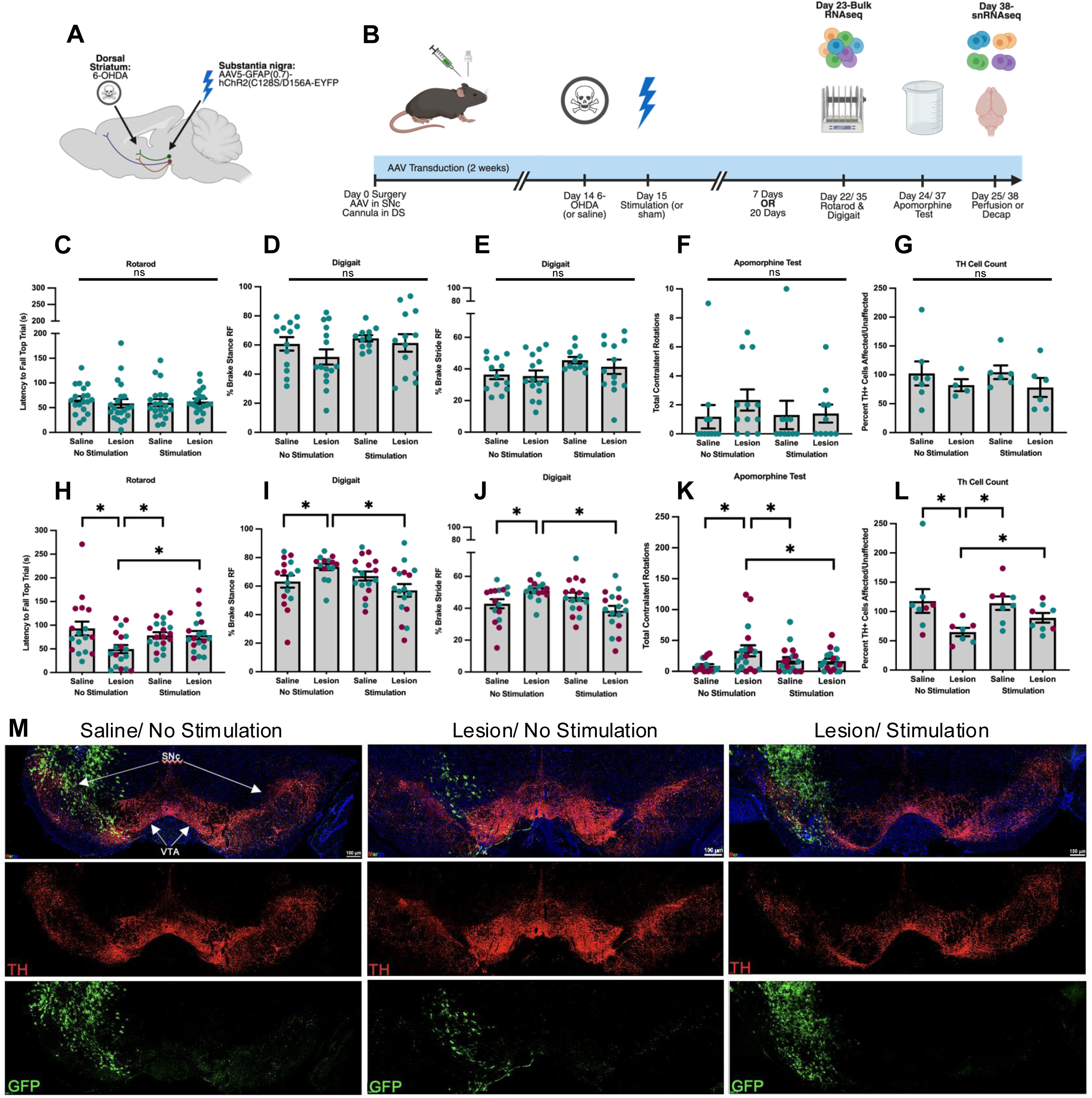
Optogenetic stimulation of astrocytes attenuates 6-OHDA induced motor deficits and Th cell death. (A) Schematic depicting the unilateral 6-OHDA lesion within the dorsal striatum and the AAV injection into the SN. (B) Experimental timeline. All rats underwent stereotaxic surgery followed by 6-OHDA (or saline) and/ or nigral astrocyte stimulation (or no stimulation control). Behavioral testing started either 1- or 3-weeks post-lesion. (C-F) There were no significant differences on any behavioral measure assessed 1-week post-lesion. (G) Cell counts comparing the number of Th+ cells in the affected over unaffected SN 1-week post-lesion. There were no significant differences between any of the groups. (H-K) At 3 weeks post-lesion, lesioned, no stimulated rats had significant motor impairments across all behavioral tests compared to saline, no stimulated controls. This was attenuated by optogenetic stimulation of nigral astrocytes; lesioned, stimulated rats performed comparably to controls. (L) Cell counts comparing the number of Th+ cells in the affected over unaffected SN 3-weeks post-lesion. There was a main effect of lesion. Simple t tests comparisons revealed the lesion/ no stimulation rats greater Th+ cell loss compared to all other groups and this was attenuated with optogenetic stimulation. (M) Representative images (20X) at 3 weeks post-lesion. GFP is shown in green, Th in red and DAPI in blue. Data are represented as mean ± SEM. Males are shown in teal, females are shown in maroon. **\****p* < 0.05.

As expected, rats assessed at 1-week post-lesion showed no deficits on the rotarod (F_1,76_=0.502, *p*=0.481), the percent brake stance of the contralateral (affected) fore paw (F_1,48_=0.343, *p*=0.561), the percent brake stride of the contralateral (affected) fore paw (F_1,48_=0.216, *p=*0.644), or the number of contralateral rotations in the apomorphine-induced rotations test (F_1,39_=0.438, *p*=0.512) (Figure 1C-F). At 3-weeks post-lesion, however, significant motor impairments were observed where lesioned rats performed worse than controls on the rotarod (planned comparison: t(33)=2.588, *p*=0.014), showed significant gait abnormalities (percent brake stance t(29)=2.0913, *p*=0.045; percent brake stride t(29)=2.4842, *p*=0.019) and greater contralateral rotations on the apomorphine-induced rotations test t(32)=2.495, *p*=0.018. Remarkably, lesioned rats that received one 30-min session of optogenetic stimulation of nigral astrocytes showed a complete attenuation of motor impairments such that they were no different from saline/ no stimulation controls) on the rotarod (F_1,69_=4.937, *p*=0.03), the percent brake stance of the contralateral (affected) fore paw (F_1,61_= 7.538, *p*=0.008), the percent brake stride of the contralateral (affected) fore paw (F_1,61_=10.610, *p*=0.002) or the number of contralateral rotations in the apomorphine-induced rotations tests (F_1,65_=4.734, *p*=0.033). This suggests a protective effect of stimulating astrocytes via optogenetics (Figure 1H-K).

### Optogenetic stimulation of SNc astrocytes attenuates dopaminergic cell loss

To confirm whether the motor deficits were related to nigral DAergic cell loss, the percentage of tyrosine hydroxylase (Th+, the rate-limiting enzyme in the biosynthesis of dopamine) cells in the affected (ipsilateral to the lesion) over number of cells in the unaffected SN was calculated to determine Th+ cell loss following 6-OHDA exposure. There was no significant loss of Th+ cells at 1-week post-lesion (F_1,19_=0.31, *p*=0.863) (Figure 1G). However, by 3-weeks the lesioned (with no stimulation) rats had a lower percentage of Th+ cells compared to all other groups (F_1,27_=3.358, *p*=0.033). Unpaired t-tests confirmed lesion/ no stimulation rats had a significantly lower percentage of Th+ cells compared to the saline/ no stimulation group t(13)=2.3289, *p*=0.037, the saline/ stimulation group t(13)=3.5174, *p*=0.004 and the lesion, stimulation group t(13)=2.2071, *p*=0.046. However, lesioned rats with stimulation did not differ significantly from saline, no stimulation controls t(14)=1.3137, *p*=0.210, suggesting that as expected, 6-OHDA induced DAergic cell death and that optogenetic stimulation of astrocytes attenuated this (Figure 1L).

### Bulk RNA-sequencing reveals changes in microglial-related genes

While the effects of optogenetics on neuronal activity, function and neuronal networks have been well described, the effects of optogenetic stimulation of astrocytes on the regional milieu are less understood.^20^ To assess the network at a time of ongoing degenerative processes (or attenuation thereof), we employed bulk RNA-seq of the SNc at one-week post-lesion/stimulation (Figure 2). We found that in the absence of any stimulation, there were 53 differentially expressed genes (DEGs) (21 upregulated and 32 downregulated) when comparing the lesioned and saline treated rats. Interestingly, this included an increase in *Th* mRNA expression, indicating potential dopaminergic expression pattern differences were induced by the lesion alone. This may reflect a potential compensatory mechanism to negate the ongoing death of dopaminergic neurons in the SN (Supplemental File 6). Indeed, increased Th expression and activity has been observed following neurotoxicant exposure.^63,64,65^

**Figure 2.**
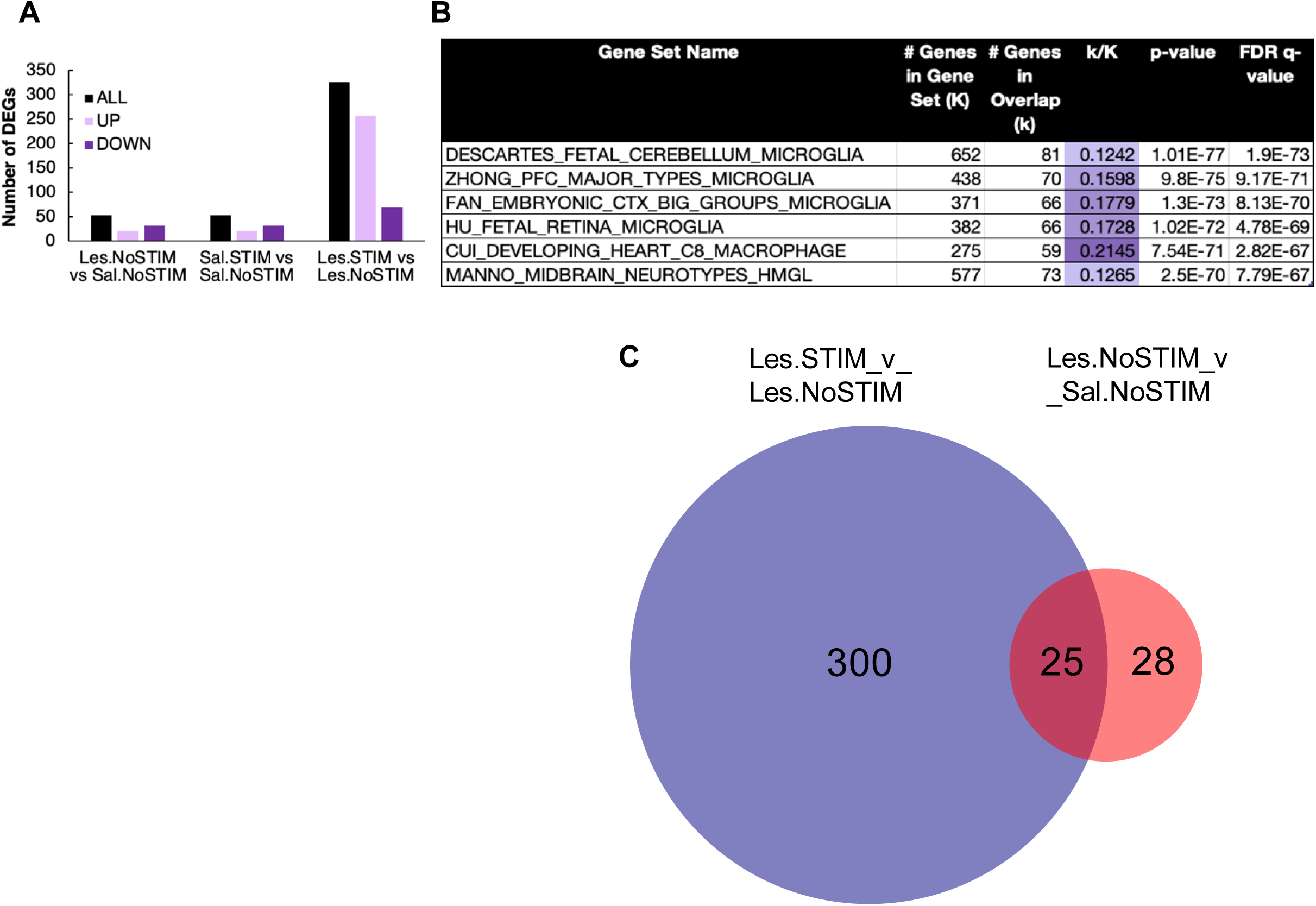
Stimulation of nigral astrocytes induces microglia activation in a 6-OHDA model of neurodegeneration. (A) Bulk RNA-seq of the SN 1-week post-lesion. There were 53 DEGs (21 upregulated and 32 downregulated) between lesion/ no stimulation rats and saline/ no stimulation rats (effect of 6-OHDA alone). There were 53 DEGs (21 upregulated and 32 downregulated) between saline/ stimulated rats and saline/ no stimulation controls (effect of optogenetic stimulation alone). Between lesioned rats with stimulation and lesioned rats without stimulation, there were 325 DEGs (256 upregulated and 69 downregulated). (B) Gene Set Enrichment Analysis (GSEA) of the 325 DEGs between lesioned rats with stimulation verses lesioned rats without stimulation revealed an enrichment in microglia-related pathways. This suggests that the neuroprotective effects of stimulating nigral astrocytes may be partly mediated through changes in microglial activity. (C) Overlap of DEGs between the lesion/ no stimulation vs. saline/ no stimulation comparison and the lesion/ stimulation vs. lesion/ no stimulation comparison. 25 DEGs appeared in both lists. 17 DEGs downregulated with 6-OHDA were upregulated with stimulation, and 2 DEGs that were upregulated with 6-OHDA and then downregulated with stimulation. Image generated using DeepVenn.

Importantly, the astrocyte optogenetic stimulation alone provoked gene expression changes, with 53 DEGs found between saline/stimulated rats and the saline/no stimulated rats (21 upregulated and 32 downregulated). Furthermore, 325 DEGs (256 upregulated and 69 downregulated) were evident when comparing lesioned rats with stimulation and lesioned rats without stimulation (Figure 2A and Supplemental File 3). Interestingly, the comparison of stimulation versus no stimulation lesion groups showed almost 5 times higher DEGs than any other comparison, potentially suggesting that these two groups have undergone a greater state of change than even the lesion/saline comparison.

To better understand functional changes within the SN at 1-week post-lesion, Gene Set Enrichment Analysis (GSEA) was employed. Some of the top pathways called between lesioned rats without stimulation and saline/no stimulation controls included a response to extracellular matrix, and a response to virus-an effect likely driven by the change in immune-related genes such as *Ifitm1, Ifit3, Cxcl10* and *Irf7*. Of particular relevance to PD, GSEA revealed that DEGs between the stimulation and no stimulation saline groups (without lesions) were enriched in pathways related to aging, secretory vesicles and cell-cell signaling (Supplemental File 6). Interestingly, several collagen-related genes, Col1a1, Col3a1 and Col1a2 were differentially expressed following astrocyte stimulation. This could suggest that under basal conditions optogenetic stimulation of astrocytes may induce changes in the extracellular matrix, at least within the substantia nigra.

Surprisingly, despite having received astrocyte simulation, GSEA analysis revealed that the top enriched pathways between lesioned rats that were either stimulated or not were actually associated with microglia and macrophage function (Figure 2B). Indeed, the stimulation increased microglia related genes such as *Cd68*, *Tyrobp*, *Csf1r*, *Cx3cr1* and *Trem2* in the lesioned rats (Supplemental File 3), suggesting a potential role for microglia involvement in the neuroprotective effects of nigral astrocyte activation.

Furthermore, there were 25 DEGs that overlapped between the effects of 6-OHDA alone and the effects of stimulation following 6-OHDA (Figure 2C). Of these 25 DEGs, 17 were decreased with 6-OHDA alone, an effect that was reversed with nigral astrocyte stimulation, suggesting that these could be causally involved in the detrimental effects of the lesion that is reversed with stimulation (Supplemental File 3). Only 2 DEGs were increased in the lesioned rats compared to controls, and then decreased in lesioned/ stimulated vs lesioned/ no stimulated rats, (*AABR07046778.1* and *Prkcd*). One of these genes, *Prkcd*, encodes for protein kinase C delta, which is increased in microglia within the ventral midbrain of patients with PD.^66^ *Prkcd* deletion dampens microglial inflammatory responses, attenuates dopamine cell death and reduces motor impairments in LPS and MPTP models of PD.^66^

### 6-OHDA induces gene expression changes in glial cell types

Although the unilateral striatal 6-OHDA lesion model has long been employed across many contexts, to our knowledge, a complete profile of cell type specific transcriptomic changes has not been conducted. Accordingly, we employed snRNA-seq on SN samples collected 3-weeks post-lesion, at a time when behavioral and cellular deficits were present. We found that 6-OHDA alone induced extensive gene expression changes across all cell types, but most notably in excitatory neurons (10117 DEGs), and interneurons (3894 DEGs) (Figure 3 and Supplemental File 4). Surprisingly, there were also substantial changes in the gene expression of oligodendrocyte lineage cells, with 4652 and 3972 DEGs in mature and immature oligodendrocytes, respectively (Figure 3 and Supplemental File 4). There were moderate changes in gene expression of Th + neurons (1561 DEGs), astrocytes (1220 DEGs) and OPCs (952). Finally, very few DEGs were observed in the microglia cluster (97), and the astrocyte/ endothelial cell (EC) cluster, which only had 9 DEGs after 6-OHDA treatment (Figure 3). Across nearly all cell clusters, there were more downregulated DEGs following 6-OHDA treatment compared to saline (Supplemental File 4). Perhaps unsurprisingly, given our 1-week post-lesion data, the reverse was true in microglia, wherein approximately 70.6% of DEGs were actually upregulated after the neurotoxin treatment (Supplemental File 4).

**Figure 3.**
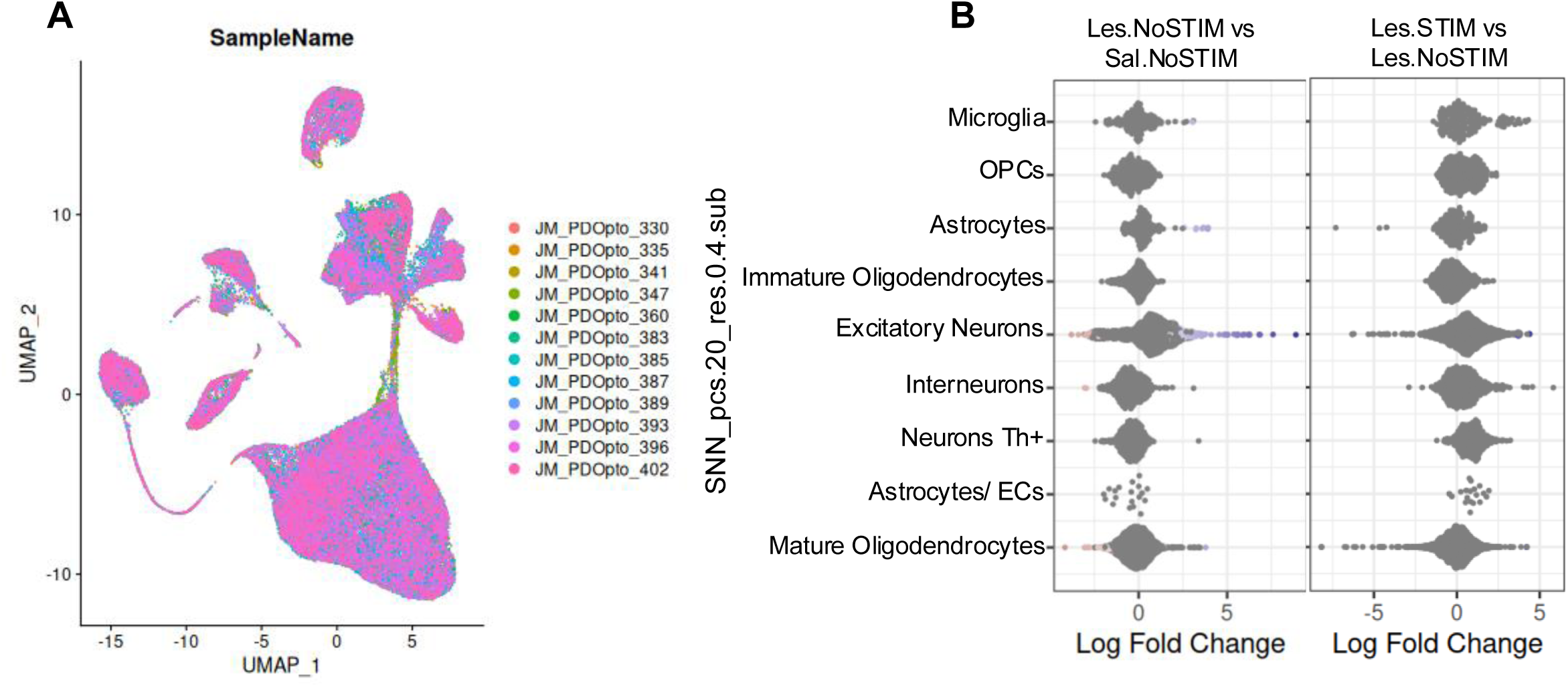
6-OHDA induces gene expression changes in all glial cell types. (A) snRNA-seq of the SN 3-weeks post-lesion. UMAP depicting independent SN samples. All samples were represented in each distinct cell cluster. Cells are colored according to independent sample. (B) Beeswarm plots showing cell abundance changes in nigral cell populations between lesion, no stimulated rats and saline, no stimulation controls and between the lesion/stimulation verses lesion/ no stimulation groups.

Using GSEA, we looked at the enriched pathways for the top 100 DEGs of each cell cluster. The terms “midbrain neurotypes,” “synaptic function”, “post-synapse”, “synapse”, and “development of neurons” were called in many cell clusters, including astrocytes, immature oligodendrocytes, interneurons and Th+ neurons (Supplemental File 7). In mature oligodendrocytes, the top enriched pathways were related to synaptic function: “synapse”, “post-synapse” and “synaptic membrane,” were in fact, all listed. In astrocytes, the DEGs were related to “post-synapse”, “embryonic cortex astrocytes” and “cell junction organization.” (Supplemental File 7). For microglia, GSEA analysis of all 97 DEGs was performed. Some of the top pathways were related to “Parkinson disease” and “cell projection organization” (Supplemental File 7). Given the low number of DEGs in the astrocyte/EC cluster, pathway enrichment analysis was not performed.

### Optogenetic stimulation of astrocytes induces long term gene expression changes in microglia

By 3-weeks following 6-OHDA administration, the motor deficits and Th+ cell loss present in the lesioned rats were significantly attenuated by the optogenetic astrocyte stimulation, therefore, snRNA-seq of the SN was conducted at this time. The snRNA-seq uncovered substantial gene expression changes in excitatory neurons (10377 DEGs) and interneurons (3194 DEGs) when comparing lesioned rats with and without stimulation (Figure 3 and Supplemental File 5). Once again, the mature and immature oligodendrocytes both had significant changes in gene expression, with 6114 and 4006 DEGs, respectively (Figure 3 and Supplemental File 5). The OPC, astrocyte and Th+ neuron clusters all had more moderate changes in gene expression (1806, 1220 and 1109 DEGs, respectively), while the astrocyte/ endothelial cell cluster had only 11 (Figure 3 and Supplemental File 5). While there were very few microglial gene expression changes with 6-OHDA alone (97), there were 1276 DEGs in microglia induced by the stimulation in the context of 6-OHDA exposure (Figure 3 and Supplemental File 5). This further supports the notion that optogenetic stimulation of astrocytes impacts the glial cell milieu in the SN when neurodegenerative processes have been triggered. We posit that it is likely that the optogenetic stimulation is neuroprotective by direct and indirect influences that astrocytes have on the DAergic cellular environment.

### Loss of Th+ neurons increases Th expression in oligodendrocytes

In parallel to our immunohistochemical findings, the number of *Th*+ neuron neighborhoods was decreased with 6-OHDA lesion, and this loss was attenuated with astrocyte stimulation (Figure 4B-D). Interestingly, *Th* expression was not only seen in neurons, as snRNA-seq data revealed a significant cluster of mature oligodendrocytes and a small cluster of astrocytes and microglia that also express *Th* (Figure 4E). Although this finding may seem remarkable, there is some limited data showing *TH* expression in oligodendrocytes and that it is altered PD patients.^47^ However, we made the very novel finding that *Th* expression levels in oligodendrocytes were significantly increased following lesion (suggesting a potential compensatory effect), but that this was attenuated with astrocyte stimulation (Figure 4E). To confirm *Th* gene expression in oligodendrocytes results in Th protein, immunohistochemical analysis was conducted. Indeed, this confirmed the presence of Th+ oligodendrocytes in the SN (Figure 4G). In addition, we confirmed that TH+ oligodendrocytes are also present in the SN of human post-mortem tissue, suggesting this is not a species-specific anomaly (Figure 4G). Like midbrain DA neurons, mature oligodendrocytes express high levels of dopa decarboxylase (DDC), the enzyme that converts the precursor, L-DOPA, into dopamine (Figure 4F). Interestingly, the number of mature oligodendrocytes expressing DDC appeared to decrease slightly with 6-OHDA treatment, and once again, this effect was attenuated with optogenetic stimulation (Figure 4F).

**Figure 4.**
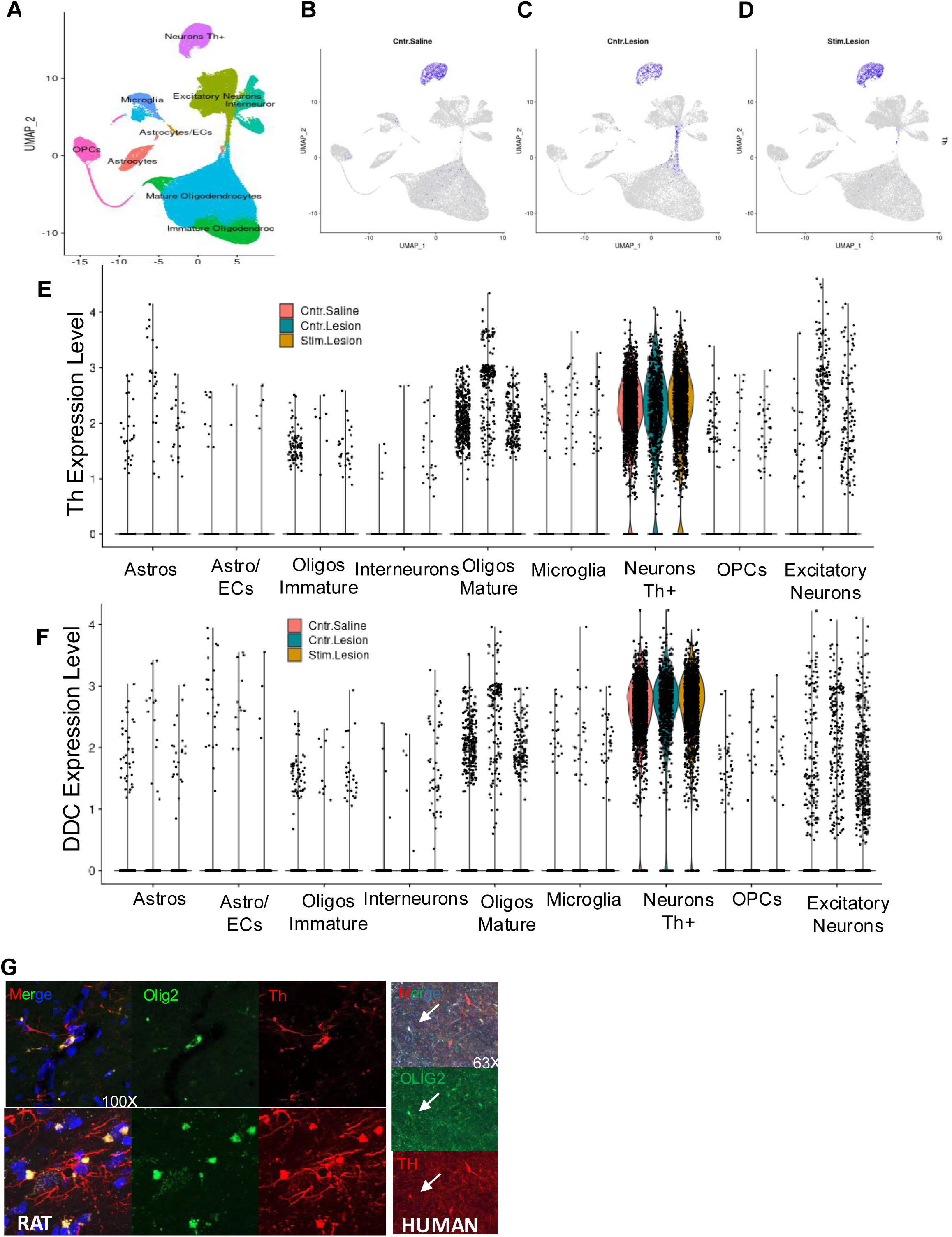
Loss of SNc dopamine neurons induces Th+ expression in oligodendrocytes. (A) snRNA-seq of the SN 3-weeks post-lesion. UMAP depicting all distinct cell clusters across the groups. Cells are colored according to cell type. (B-D) UMAP highlighting Th+ cells in the saline/ no stimulation (Cntr.Saline) (B), lesion/ no stimulation (Cntr.Lesion) (C) and lesion/ stimulation (Stim.Lesion) groups (D). (E) Violin plots showing Th expression in each cell type across saline/ no stimulation, lesion/ no stimulation and lesion/ stimulation groups. (F) Violin plot showing DDC expression in each cell type across saline/ no stimulation, lesion/ no stimulation and lesion/ stimulation groups. (G) Representative images (100X) showing Th+ oligodendrocytes in the rat SNc. Representative image (63X) showing TH+ oligodendrocytes in the human SNc of 37-year-old lymphoma patient.

## Discussion

In the current work, we show that a single session of optogenetic stimulation of nigral astrocytes, shortly after exposure to 6-OHDA, is sufficient to reduce motor impairment and attenuate dopaminergic cell loss. This builds on previous work demonstrating *in vitro* optogenetic stimulation of astrocytes prevented the neurodegenerative effects of the neurotoxin MPTP and that *in vivo* stimulation promoted astrocytic growth factor, FGF2, and induced the functional repair and integration of stem cells.^61^ Here, rather surprisingly, bulk RNA-sequencing revealed that these neuroprotective effects appeared to be partially mediated through changes in microglial activity. Growth factors, cytokines, gliotransmitters, mitogenic factors and even extracellular vesicles represent some of the numerous secreted mediators that facilitate astrocyte-microglia crosstalk.^67^ Given that nigral astrocyte stimulation occurred shortly after 6-OHDA exposure, the astrocytic release of anti-inflammatory chemokines and cytokines, as well as growth factors, might represent some of the early released mediators that influenced microglial response to ongoing neurotoxic cascades. Hence, assessing the astrocyte transcriptome immediately or very shortly after optogenetic stimulation would provide insight into which astrocytic pathway(s) are mediating the subsequent alterations in microglia.

It is particularly important for future studies to address whether astrocyte-microglia crosstalk is necessary for the neuroprotective effects of optogenetic stimulated nigral astrocytes. To address microglial involvement one could use the approach taken by several studies that have used CSF1R inhibitors, a crucial growth factor receptor required for microglial survival, to deplete microglia within the SNc.^68,69,70,71,72^ While the loss of nigral microglia improved motor impairments and dopamine cell death in some instances,^68,69^ in other cases, the loss of microglia greatly exacerbated the deficits induced by the neurotoxins MPTP^70,71^ or 6-OHDA.^13^ Hence, the role of microglia cannot be simply reduced to one of “protective vs toxic”. Rather, there are a host of intermediate to extreme classical and alternative activation states that characterize the response adopted by these cells with their detection of insults in their microenvironment. It is also important to note that the timing of these interactions with relation to neuronal cell death may be critical to the effects observed. Nevertheless, these findings (along with work from the Liddelow lab),^24,38,73^ highlight both the complex and integral role of microglia in modulating dopamine neuronal survival, and given their extensive crosstalk with astrocytes, may position microglia as an important intermediary determinant between nigral astrocyte optogenetic stimulation and neuroprotection.

snRNA-sequencing and immunohistochemistry revealed that besides dopaminergic neurons, Th, the rate-limiting enzyme in dopamine synthesis, is expressed in other cell types including astrocytes, and most notably, mature oligodendrocytes. Though not commonly acknowledged, TH is indeed expressed to a certain degree in oligodendrocytes and was reported to be elevated in the PD brain^47^. Intriguingly, *Th* expression in mature oligodendrocytes increased following 6-OHDA administration, but was attenuated with optogenetic stimulation of nigral astrocytes. This could reflect a compensatory mechanism, but although speculative, might also stem from changes in the myelination of neuronal projections that occurs in PD.^74,75,76,77,78^ Like all glial subtypes, the notion that oligodendrocytes may be critically involved in PD is not novel, indeed, a 2020 paper that integrated numerous genome-wide association studies and single-cell transcriptomic data sets found that besides monoaminergic neuron populations, oligodendrocytes are also genetically associated with PD.^79^ Likewise, Webber and colleagues found an association between PD-risk genes and oligodendrocytes within the human SN that was driven by a fraction of risk genes distinct from those that linked dopaminergic neurons to PD risk,^80^ suggesting that oligodendrocyte-driven susceptibility to PD may represent a unique pathogenic pathway.

Extensive changes in oligodendrocyte gene expression have been noted in PD, particularly within the SN^47,50^ but also in the caudate nucleus and putamen^81^ and even cortical areas.^82^ Changes in cytoskeleton organization/ dynamics,^83,84^ cellular stress response,^83^ autophagy and chaperone-mediated protein folding^85,86^ have all been identified as pathways enriched in oligodendrocytes in patients with PD. A subpopulation of oligodendrocytes identified in PD was also enriched in interferon response, negative modulation of the immune system and oxidative phosphorylation.^85,86^ This suggests important inflammatory changes in oligodendrocytes in PD.^85^ Whether these transcriptomic shifts in oligodendrocytes are a major determinant of PD pathogenesis or a secondary effect to disease progression remains to be elucidated, nonetheless, it is evident that oligodendrocytes are more critically involved in PD than previously believed.

Glial-glial interactions are important modulators of neuroinflammation, a key feature of PD, and driving force of neurodegeneration. For instance, pro-inflammatory cytokine release from activated microglia can induce a neurotoxic phenotype of reactive astrocytes, characterized by increased oxidative stress and reduced neurotrophic release,^19^ which could perpetuate further neuroinflammation. Similarly, astrocytic derived ATP can induce release of the pro-inflammatory cytokine, IL-1β, from microglia,^87^ whereas astrocytic TGF-β can influence microglial morphological state and funcitoning.^88^ Though known for their role in myelination, oligodendrocytes are also modulators of neuroinflammation. In fact, both astrocytes and oligodendrocytes can influence innate and adaptative neuroimmune functions through the release a variety of growth factors, interferons and cytokines such as, but not limited to, FGF2, IFN-γ, IL-1β and IL-17.^89^ Outcomes between this crosstalk of astrocytes and oligodendrocytes can be beneficial or detrimental, but importantly, helps to modulate the inflammatory environment.^89^ For instance, activated astrocytes can induce apoptosis of oligodendrocytes via TNF signaling or recruit oligodendrocytes to sites of inflammation^90^ whereas CCL2 release from oligodendrocytes reduces IL-6 signaling in astrocytes, also decreasing inflammation.^89,91,92^

Despite directly stimulating nigral astrocytes in the current work, extensive gene expression changes were observed in microglia and oligodendrocytes, including many inflammatory and immune-related genes. Therefore, optogenetic stimulation of astrocytes may impact neurodegeneration and neuroinflammation via altered glial-glial interactions/ crosstalk, though additional experiments would be necessary to confirm this.

### Limitations of the study

This study has several limitations. Firstly, male and female rats were only used in behavioral testing at the later time point (3-weeks post-lesion). Although no sex differences were observed in this experiment, it is possible that sex differences in early processes may differ.^93,94,95^, Secondly, although 6-OHDA is a well-established model of neurodegeneration, it fails to recapitulate critical aspects of PD, thus limiting the generalizability of results to PD. Investigating optogenetic stimulation of astrocytes in more etiologically relevant models such as the injection of α-synuclein preformed fibrils into the brain or the overexpression of human α-synuclein with a PD-associated A53T mutation may help to test the therapeutic role of astrocytes in PD.

In conclusion, we found that a session of optogenetic stimulation of nigral astrocytes is sufficient to attenuate motor deficits and dopamine cell death induced by the neurotoxin, 6-OHDA. We also show that these neuroprotective effects are associated with glial-glial interactions both at 1- and 3-weeks post-lesion. This study clearly demonstrates the importance of crosstalk between astrocytes, microglia and oligodendrocytes in modulating outcomes of nigral dopamine cell degeneration relevant for PD.

## Methods

### Animals

A total of 118 adult male and 36 adult female Long Evans rats (8-10 weeks) were obtained from Charles River Laboratories, Québec. All rats were individually housed in standard Green line IVC cages (39.5 x 34.6 x 21.3 cm) with basic enrichment and maintained on a 12-h light/dark cycle with lights on at 07:00 hours. *Ad libitum* access of standard chow and water was provided with room temperature maintained at ∼21°C. The endpoint for experimental rats was reached if they lost > 15% of their body weight. All animal procedures including euthanasia were approved by the Carleton University Committee for Animal Care in accordance with the standards and guidelines established by the Canadian Council for the Use and Care of Animals in Research.

### Stereotaxic Surgery

While under isoflurane anesthetic (Fresenius Kabi), all rats had 1.0 μ1 of AAV5-GFAP(0.7)-hChR2(C128S/D156A)-EYFP (1.4 × 10^13^ GC/ml) (Vector Biolabs) infused unilaterally into the SNc at a rate of 0.1 μ1/min for a total of 10 mins, followed by a 10 mins period where the injector was left undisturbed to allow for diffusion away from the injector tip. To facilitate easy delivery of light to the virally-transduced astrocytes of the SNc, an optical fibre implant was placed at the same Bregma coordinates (Males: AP: −5.8 mm; ML: 2.0 mm; DV: −7.2 mm; Females: AP: −5.8 mm; ML: 2.0 mm; DV: −7.1 mm) and secured using dental cement (Stoelting-51458) and surgical screws (McMaster-Carr). Optical implants were made by placing 300 μm Core Multimode optical fiber (Thorlabs) into a stainless alloy ferrule drilled to 350 μm (Fiber Instrument Sales Inc.) and secured using heat curable epoxy resin (Precision Fiber Products Inc). Optical implants were polished using 5 μm silicon carbide polishing paper, 3 μm and 1 μm aluminum oxide polishing paper and 0.3 μm calcined alumina polishing paper (Thorlabs) before use. A guide cannula (22 gauge) (Protech International Inc.) was subsequently implanted into the dorsal striatum (Bregma coordinates: Males: AP: 1.2 mm; ML: 3.0 mm; DV: −5.4 mm; Females: AP: 1.2 mm; ML: 3.0 mm; DV: −5.3 mm) and also secured using dental cement and surgical screws.

### 6-hydroxydopamine Lesion

Two weeks after the stereotaxic surgery, rats were subjected to a unilateral 6-OHDA lesion (or saline control) that was ipsilateral to the AAV-injected SN. 6-OHDA (Sigma Aldrich H4381) was dissolved in a solution of 0.02% ascorbic acid in 0.9% saline. Under isoflurane anesthetic, rats were injected with a 10 μg/μl solution of 6-OHDA into the dorsal striatum. This was done at a rate of 0.4 μl/min over 5 mins for a total infusion of 2.0 μl. The injector remained in place for 1 min following the infusion. Control groups received 2.0 μl of saline injected at the same rate.

### Optogenetic Stimulation of Astrocytes

One day after the 6-OHDA (or saline) injection, rats underwent optogenetic stimulation of astrocytes in the SNc. Rats were connected to the LPS-0473 DPSS Laser System (Laserglow Technologies) via the optical implant embedded into their SNc. The single optogenetic stimulation session began with the laser light on for 5 mins, pulsing at a frequency of 5 Hz (B&K Precision), followed immediately by a 5 mins period in which the laser light was off. This cycle repeated 3 times for a total of 30 mins. Control rats (“no stimulation”) were connected to the laser but the light remained off for the full 30 mins period.

### Motor Behavior Analysis

At 1-week (males only) or 3-weeks post-lesion (males and females), groups underwent a battery of behavioral tests to assess potential changes in motor behaviors, including the rotarod, gait analysis and apomorphine-induced rotations test.

### Rotarod

At 1-week days or 3-weeks post-lesion, rats underwent the accelerating rotarod (San Diego Instruments) test to assess potential changes in motor learning and coordination. Rats underwent 4 separate trials of up to 300 sec in which the speed increased slowly from 0-40 rpm. Trials were separated by at least 30 mins to prevent fatigue. The latency to fall off (s) was recorded for each trial.

### DigiGait

Immediately following the rotarod (ie. same day of testing - refer to Figure 1B), the DigiGait (Mouse Specifics, Inc.) apparatus was used to detect any changes in gait. Rats walked on the DigiGait treadmill at a rate of 21 cm/s at a 0° incline for approximately 3 sec. Several indices of gait such as the percent brake stance and percent brake stride were recorded. Rats that did not walk on the treadmill for at least 3 sec were excluded from the analyses (Supplemental File 1).

### Apomorphine-Induced Rotations Test

Two days following the rotarod and DigiGait testing (on day 24 or 37), rats underwent the apomorphine-induced rotations test. A 0.5 mg/kg subcutaneous dose of apomorphine (Sigma Aldrich A4393-100MG) was administered 5 mins prior to testing which lasted for 25 mins in a clear glass cylinder. All tests were recorded and later scored by a rater. The number of both contralateral and ipsilateral rotations were counted. A rotation was counted if the rat made a continuous, uninterrupted turn (360^°^) with both its head and body in one direction.

### Immunohistochemical Analysis

For immunohistochemical analysis, rats were injected with Euthanyl and then perfused intracardially with PBS followed by fixation using 4% paraformaldehyde. Whole brains were extracted and post-fixed for 24 hrs in 4% paraformaldehyde, followed by 72 hrs in a 30% sucrose solution. Fixed brains were wrapped in aluminum foil and stored at −80°C until further processing. Brains were sectioned at 15 μm. For all immunohistochemical analyses, sections were first incubated in pre-block (10% horse serum in PBS-Triton) for 1 hr. Primary antibodies were applied overnight at room temperature. The following day, sections were washed in PBS and then secondary antibodies were applied for 2 hrs at room temperature. Subsequent PBS washes were applied and then tissue was mounted using hardset DAPI (Vectashield). A list of all antibodies used and their respective dilutions can be found in Table 1.

### TH-positive Cell Counting

Dopaminergic cell loss was determined as a percentage of Th+ cells in the affected (lesioned) vs. unaffected SN for each rat. Using the Zen 2.6 system, Th+ cells were counted first in the lesioned (or saline-injected) SN followed by the unaffected (control) SN. The number of Th+ cells in the affected side was divided by the number of Th+ cells in the unaffected side to determine the percentage. Th+ cells were counted when the Th marker overlapped with DAPI.

### RNA-sequencing analysis

For both bulk RNA-sequencing (RNA-seq) and single-nuclei RNA-sequencing (snRNA-seq) analyses, brains were collected via rapid decapitation. The substantia nigra (SN) for each sample was dissected out on ice and immediately flash frozen using dry ice. Tissue was kept at −80°C until further processing.

### Bulk RNA-seq

For bulk RNA-sequencing, the Qiagen RNeasy^®^ Midi Kit was utilized according to commercial guidelines. Briefly, 2 ml of Buffer RLT/ ®-mercaptoethanol was added to the tissue. Using a rotor, tissue was homogenized for 45 sec followed by 10 mins in the centrifuge at 3345 g for 10 mins. The supernatant was transferred out without disrupting the pellet. Next, 2.0 ml of 70% ethanol was added to the homogenized lysate and was mixed vigorously. The samples were placed in midi columns and centrifuged at 3345 g for 5 mins, and any flow through was discarded. 4 ml of Buffer RW1 was added to the column and again, samples were centrifuged at 3345 g for 5 mins. Flow through was discarded. This was repeated using 2.5 ml of Buffer RPE, followed by centrifugation for 2 mins at 3345 g. A second volume of 2.5 ml of Buffer RPE was added to the column, and samples were centrifuged for 5 mins at 3345 g. The sample was then transferred from the spin column to a collection tube where 150 μl of RNase-free water was added. Samples remained undisturbed on ice for 1 min, and were then centrifuged at 3345 g for 5 mins. A second elution was also collected.

Samples were sent to the Yale Center for Genome Analysis (New Haven, Connecticut) for whole-tissue RNA-sequencing. Quality control of samples was assessed using Agilent Bioanalyzer Gel; only mRNA with RNA integrity values (RINs) of ≥ 7.0 were used for further processing. Purified mRNA was generated using oligo-dT beads. Random primers were used for first-strand synthesis followed by a second-strand synthesis with dUTP to acquire a strand-specific library. Reverse transcription converted purified RNA into cDNA. cDNA libraries were amplified for subsequent sequencing using 75bp paired-end sequencing on an Illumina HiSeq 2500 (according to Illumina protocols). Statistical analysis was conducted similar to previous work in the lab.^96,97^ Briefly, fastq files were mapped to the Ensembl (release 95) Rattus Norvegicus reference genome (Rattus_norvegicus.Rnor_6.0.dna.toplevel) and the Rnor_6.0 reference transcriptome, using the HISAT2 v2.1.0 mapper [https://doi.org/10.1038/s41587-019-0201-4] with parameters --dta -t -k 1. The resulting bam files were quantified using StringTie v2.1.4 [https://doi.org/10.1038/nbt.3122] with default parameters. We used fastqc [https://www.bioinformatics.babraham.ac.uk/projects/fastqc/] and Qualimap [https://doi.org/10.1093/bioinformatics/btv566] for quality control of, respectively, fastq and bam files. We then used multiQC to assemble overall qc reports. Based on qc metrics and PCA, we discarded two samples and retained the remaining 17 samples for analysis. The counts files were then processed with an in-house script in R for differential expression analysis using edgeR [https://doi.org/10.1093/bioinformatics/btp616]. Gene level filtering was done using the egdeR function filterByExpr requiring min.count=10 and min.prop=0.7, providing the experimental groups structure. Counts were then processed using cqn [https://doi.org/10.1093/biostatistics/kxr054] to create an offset matrix to be used for GC correction as input to edgeR to test for differential expression. Nominal p-values were FDR corrected and an adjusted p-value < 0.05 was considered significant.

### Sample preparation for nuclei isolation

snRNA-seq samples were prepared and sequenced at the Princess Margaret Genomics Centre (Toronto, Ontario) using the 10X Genomics Chromium platform. Briefly, nuclei isolation consisted of weighing and cutting the tissue and then transferring to 1 ml lysis buffer for 5 mins. Using a Rainin wide bore pipette, samples were then transferred to, and homogenized, in a glass douncer (KIMBLE Dounce tissue grinder set). A pipette (P1000 wide bore) was used to mix the samples, and then the samples were placed in a 2 ml low bind tube (Eppendorf). Samples were then spun at 800g for 10 mins while maintained at 4°C. The supernatant was removed, and the pellet resuspended in 2 ml of wash buffer. The mixture was pipetted 20 times, and the lysate was filtered through a 30 μm Miltenyi filter and collected into another 2 ml tube (Eppendorf). The samples were spun once again at 800g at 4°C for 10 mins. The supernatant was removed, and 2 ml of wash buffer added to resuspend the pellet. Samples were centrifuged at 800g for 10 mins at 4°C. The supernatant was removed, the pellet resuspended in 1 ml of wash buffer and then filtered with a 40 μm Flowmi cell strainer and transferred to 1.5 ml low bind tube (Eppendorf) on ice. Nuclei counts were done on a disposable cell counter and fluorescent microscope using a mix of 10 1l of sample and 10 μl AOPI (Acridine Orange-Propidium Iodide).

### snRNA-seq

Sequencing reads were pre-processed using CellRanger (v7.0.0; 10X Genomics Inc.) using the mRatBN7-2-2024-A reference transcriptome. We sequenced an estimated 10,000-15,000 single-cells per sample, resulting in ∼6,000 usable cells per sample, and an average depth of about 50,000 reads per cell.

We processed the data using R (v4.3.2) and Seurat (v4.3.0.1) [https://doi.org/10.1016/j.cell.2021.04.048]. For each sample, cells with less than 1,500 reads or 1,000 expressed genes were removed. We then used the MiQC package (v1.8.0) [https://doi.org/10.1371/journal.pcbi.1009290] with default parameters to remove cells with abnormal mitochondrial content, and the scDblFinder package (v1.14.0) [https://doi.org/10.12688/f1000research.73600.2] with default parameters and the estimated doublet rate from the 10X Genomics website (https://kb.10xgenomics.com/hc/en-us/articles/360001378811-What-is-the-maximum-number-of-cells-that-can-be-profiled) to infer and remove doublets/multiples. We SCT-transformed the expression levels and identified the top 2000 variable features using the Seurat SCTransform function method=“glmGamPoi”, vst.flavor=‘‘v2, after removing noise genes. We then integrated the samples after identifying 2000 integration anchors retaining the top 20 principal components (PCA). The UMAP reductions were computed from the integrated dataset. The Seurat FindNeighbors function was used to build a neighborhood graph and the FindClusters function was used to identify clusters with a resolution range from 0.2 to 1.8. We then estimated marker genes by differential expression in a one vs all setting using the wilcoxauc function of the Presto package (v1.0.0). We selected 0.4 as final resolution, resulting in 17 clusters, based on known marker gene segregation as well as marker gene analysis. Cluster 9 was further split in 3 subclusters, for a final total number of 19 clusters. Marker gene analysis was rerun on the 19 clusters, which were annotated by using a combination of known marker genes and manual annotation. Clusters with similar annotations were then merged.

#### Differential Abundance

Differential abundance analysis was performed using the MiloR package [https://doi.org/10.1038/s41587-021-01033-z] with default parameters, following the recommended steps. We converted the Seurat integrated object to a Milo object, retaining the computed PCA and UMAP. The kNN-graph was computed from the Seurat object graph adjacency matrix with k=60 (i.e. 5*N of samples as recommended). We computed the cell neighborhoods using prop=0.1, k=60, and the first 20 PCs. The number of cells in each neighborhood was counted and a neighborhood distance matrix estimated. Finally, we computed the neighborhood graph. We then inferred differentially abundant (DA) neighborhoods by defining a design matrix accounting for individual pairing of samples, and using the algorithm’s fdr.weighting scheme by the PCA based graph-overlap. We then tested the neighborhoods for differential abundance considering two contrasts, Stimulated vs Controls within the Lesion group and Lesion vs Saline within the Control group. We did not find significant changes at spatialFDR <0.05, although many neighborhoods underwent strong shifts in number. We annotated the neighborhoods with the cell type annotation from the established annotation and clustered together any DA neighborhood with the same cell type annotation and while retaining the direction of change. This is because we hypothesized that DA neighborhoods changing in opposite directions may underlie different cell subtypes. We then computed the DA neighborhoods’ gene markers using the MiloR function findNhoodGroupMarkers, with the following parameters: assay=“counts”, aggregate.samples=true, gene.offset=true, subset.nhoods=subnhoods, where subnhoods is a vector pointing to the DA neighborhoods.

#### Cell Type Specific Differential Expression

Differential expression analysis was performed using the DEsingle package [https://doi.org/10.1093/bioinformatics/bty332] with default parameters, following the recommended steps. Briefly, the Seurat integrated object was converted to a single cell object. Outlier cells were removed. Differential expression for each contrast of interest and each cell type was then performed using the DEsingle function with default parameters and parallel=T. Differences were considered significant at FDR adjusted p-value < 0.05.

### Statistical Analyses

A 2 (lesion; saline vs. 6-OHDA) X 2 (light therapy; no stimulation vs. stimulation) analysis of variance (ANOVA) was conducted for all motor behavioral tests and cell counts using IBM SPSS Statistics (Version 26.0). When appropriate, two-tailed independent t-tests were conducted to compare individual group means (GraphPad). Statistical significance was considered when *p* < 0.05. Data are graphically represented as means ± SEM, individual data points represent one biological replicate.

## Resource Availability

**N/A**

## Lead Contact

Any requests for additional information should be directed to the lead contact, Dr. Natalina Salmaso.

## Acknowledgements

This work was supported by Parkinson Canada and by a Canada Research Chair to N.S and a Parkinson Research Consortium fellowship awarded to J.M. We would like to thank Kyle Farmer for providing experimental protocols. We would also like to acknowledge the Animal Behaviour and Physiology Core at the University of Ottawa (RRID: SCR_022882) for their help in conducting all the behavioral testing. We would like to thank the anonymous donors for their kind donation and invaluable contribution to research.

## Author Contributions

Conceptualization, formal analysis, investigation, methodology, visualization, validation, writing – original draft, J.L.M.; conceptualization, investigation, writing– review & editing, I.T.P.; investigation, visualization, validation, writing – review & editing, S.S.; investigation, visualization, validation, writing – review & editing, C.G.; conceptualization, writing– review & editing, C.A.R.; conceptualization, writing– review & editing, Shawn H.; conceptualization, writing – review & editing, M.F.; conceptualization, methodology, writing – review & editing, J.T.; conceptualization, methodology, writing – review & editing, M.S.; data curation, formal analysis, methodology, resources, visualization, validation, writing – original draft, G.C.; conceptualization, formal analysis, funding acquisition, methodology, resources, supervision, visualization, writing – original draft, N.S.

---

**Supplementary Figure 1.**
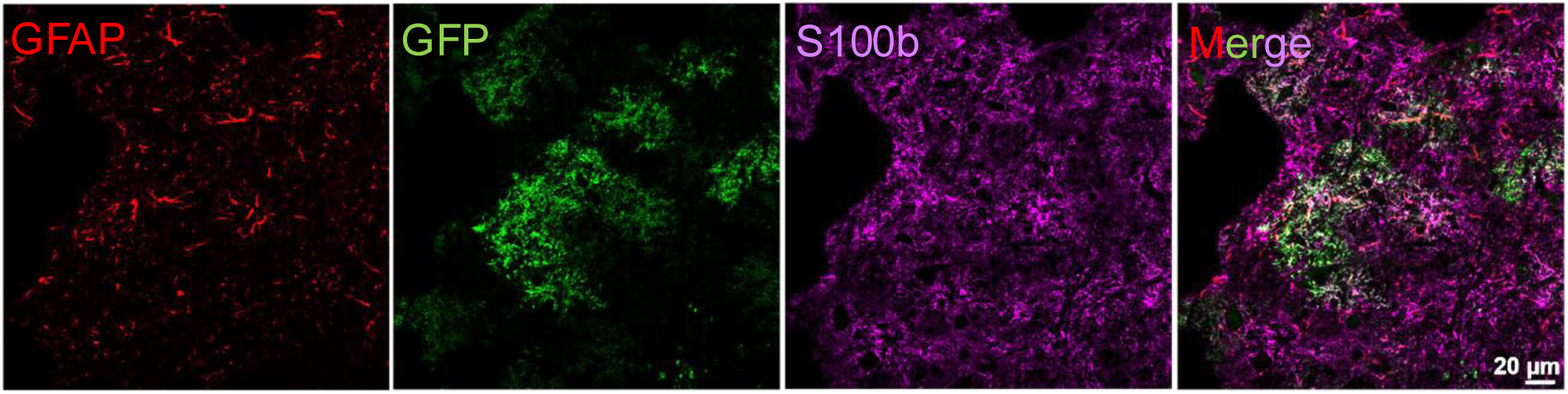
AAV expression is specific to astrocytes. A representative 63X image of a saline/ no stimulation control rat. Virally-transduced cells are GFP+ve cells and showed clear astrocyte morphology. Furthermore, GFP+ve cells co-localized with astrocyte markers such as GFAP and S100b.

